# Exogenous selenium supplements reduce cadmium accumulation and restores micronutrient content in rice grains

**DOI:** 10.1101/2024.10.16.618625

**Authors:** Falguni Barman, Titir Guha, Rita Kundu

**Affiliations:** Department of Botany, Centre of Advanced Study, University of Calcutta, 35, Ballygunge Circular Road, Kolkata-700019, India

**Keywords:** Cadmium stress alleviation, Selenium, Rice, Stress response, Micronutrients, Sulfur metabolism

## Abstract

**Purpose:** Cadmium (Cd) contamination in agricultural soil and accumulation in rice poses serious threat to human health. It is reported that Selenium (Se) can mitigate the toxic effect of Cd in rice. But the underlying mechanism of Se preventing the Cd accumulation and restoring the micronutrient content in rice grains have not been studied before. Therefore, our main aim is to reduce Cd content and restore micronutrient content in rice grain and study the mechanism.

**Methods:** Two indigenous rice genotypes (Maharaj and Jamini) were exposed to 10 and 50 µM Cd in presence and absence of Se (5µM) with a control set and assessed for plant growth, biomass, Cd content, ROS and antioxidants for Cd induced toxicity and amelioration. Genes for micronutrient transporters were studied by RT-PCR. Grain Cd and micronutrient content and agronomic parameters were also studied.

**Results:** Se supplementation increased plant growth, biomass, and yield under Cd stress. SEM and EDX analysis revealed that Se-Cd complex formed on root surfaces restricted Cd uptake by the roots preventing root damage. Soil analysis confirmed that Se decreased Cd bioavailability, restricted root to shoot Cd translocation, ultimately reducing Cd accumulation and restoring micronutrients in grain. This was further validated by fluorescent Leadmium dye staining. In (Se+Cd) treated seedlings, up-regulation of S metabolism and nutrient transporter genes also contributed to the mitigation of Cd stress.

**Conclusions:** So, the Se supplementation can be considered as a cost-effective, ecofriendly and sustainable approach to produce Cd free rice cultivation in Cd polluted soil.

## 1. Introduction

India is the second highest rice producer in world, sharing 23 percent of total rice production (120 million tons out of 503MT in 2020/21) (Pathak et al. 2020). But, in recent times contamination of heavy metals (HM) in agricultural soil declines rice productivity. Additionally, when crops are grown in heavy metal contaminated fields, harmful metals tend to accumulate in the plant tissues, posing threat to food security and human health (Zhou et al. 2015). Inhibiting this heavy metal uptake potential of the crops is a crucial objective to ensure sustainable and cleaner food production (Guo et al. 2016). Cadmium (Cd) is one of the toxic hazardous heavy metal, contaminate the soil mainly through the excessive use of phosphate fertilizers and other anthropogenic activities such as, un-regulated use of agrochemicals and sewage sludge in fields, and through mining and industrial activities (Ismael et al. 2019). Due to the increasing environmental stress and soil pollution, sustainable food production is a huge challenge (El-Ramady et al. 2019). Based on current industrial developmental trends, agricultural farm soils will face significant Cd pollution by the year 2030 and average soil Cd will range around 0.35 mg kg^-1^ which is 1.75 times higher than that of 2003 (Salmanzadeh et al. 2017).

Cd is highly mobile in nature and can be easily taken up by root and translocated to shoot. Previously, Xie et al. (2015) have been reported that rice tends to accumulate high levels of Cd, especially when Cd is present at low concentrations in soil. Cd, is a chronic carcinogen and ranked 7^th^ among the top 20 hazardous metals by the Agency for Toxic Substances and Disease Registry (ATSDR), 2012 (Nazar et al. 2012). So, Cd uptake through the contaminated rice consumption can cause multiple health hazards like diarrhea, osteomalacia, kidney diseases, even infertility (Aziz et al. 2015) in human. Additionally, Cd also restricts the nutrient uptake from soil through roots. As there are no dedicated transporters to transport Cd from soil to plants (as Cd is a non-essential metal), it mainly shares the cation transporters and transported through other mineral nutrient transporter families like IRT (Iron regulated transporter proteins), ZIP (Zinc Importer Proteins), COPT3 (Ctr/COPT-type Cu transporter proteins), NRAMP (Natural resistance-associated macrophage protein), YSL (Yellow Stripe-Like proteins) (Gao et al. 2016). This often leads to Cd accumulation as well as nutrient deficiency in rice grains (Khaliq et al. 2019).

The projected global population will increase from 7.7 billion to 9.6 billion in 2050 (Doelman et al. 2019), with this the chance of malnutrition and hunger is imminent, as the resources are limiting. This increase will occur mostly in developing countries, where chances of micronutrient deficiency and related morbidity and mortality are higher (Bouis and Welch 2010). Dietary micronutrient deficiency or “hidden hunger” affects mainly pregnant women (38%) and growing children (43%), the most vulnerable group in the population (Shahbaz et al. 2020).

Previously, several studies have been reported to reduce Cd content in rice using salicylic acid, proline, hydrogen sulfide, silicon, and nitric oxide (Rizwan et al. 2016). Along with this, exogenous application of Selenium (Se) is one of the cost-effective, eco-friendly, and sustainable method (Gao et al. 2018). Selenium (Se) is a beneficial micronutrient that reduces cancer risk and boosts immunity in animals and humans (Sonkusre 2020). Se deficiency increases the risk of susceptibility to viral infections including COVID-19 and HIV (Zhang et al. 2020a), and other diseases caused by coxsackievirus B3 and influenza A virus (Zhang et al. 2020b). Selenium (Se) (the naturally occurring amino acid) acts through the selenoproteins, containing selenomethionine, selenocysteine, and methyl-selenocysteine, diminishes the deleterious effect of several abiotic environmental stress including Cd stress (Shahid et al. 2018). In plants, Se could inhibit Cd uptake by the formation of an immobilized Se-Cd complex (Wan et al. 2016). Additionally, selenoproteins can enhance free radical scavenging activity, antioxidant property, glutathione peroxidase, and thioredoxin reductase activity (Hasanuzzaman et al. 2011). As Se and sulfur (S) are chemically similar, S metabolism can be induced by the addition of Se in plants (Chauhan et al. 2019). Moreover, S containing thiol compounds are also improved by the supplementation of Se. Thiol compounds are characterized by the presence of a sulfhydryl (−SH) functional group (Pfaff et al. 2020). They are ‘soft’ electron donors, and easily form complexes with ions of several heavy metals such as lead, mercury, cadmium (Crichton et al. 2016). Thiols form a network with the combination of sulfur-containing molecules which enhance plant stress tolerance (Zagorchev et al. 2013). The S-containing thiol compounds are called sulfur-containing defense compounds (SDCs) such as glutathione, glucosinolates, thionin and S-rich proteins can evoke “sulfur-induced resistance” (SIR) mechanism in plants. This SIR mechanism protects the plant from abiotic as well as biotic stress such as fungal pathogen and disease. Furthermore, it initiates various signal transduction pathways that are interconnected with phytohormone and ROS mediated defense system (Künstler et al. 2020). Recently (Farooq et al. 2022) and (Yang et al. 2022) reported that Se mitigate Cd toxicity by altering the thiol compounds, subcellular distribution of Cd in *Amaranthus* and morpho-physiological changes in rice respectively. But no such information is available on Se mediated up-regulation of Sulfur metabolism in Cd stressed rice, and suppression of Cd uptake by root and micronutrient enrichment in rice grain. Therefore, the main objectives of this work are to i) develop a sustainable strategy to reduce grain Cd content and ensure food safety and ii) restore micronutrients (Fe, Zn, Mn, Cu) in rice grain for dietary adequacy.

The protective role of Se against Cd toxicity was evaluated by morpho-anatomical characters (SEM images, detection of *in situ* Cd localization by fluorescence microscopy using Leadmium dye), biochemical, and molecular parameters. Pot experiment was conducted to evaluate micronutrient status in grains and to assess agronomic parameters and yield components for sustainable practice. Besides this, bioavailable Cd in soil, and human health risk were also assessed.

## 2. Material and methods

### 2.1. Plant materials and treatment condition

Two rice genotypes Maharaj (tolerant) and Jamini (sensitive) were selected from our previous study based on Cd tolerance. Rice seeds were surface-sterilized by 0.2% Dithane M-45(anti-fungal agent) and then rinsed with distilled water for 5-6 times. After that, the seeds were soaked in distilled water overnight and plated for germination. After 3 days, uniformly germinated seedlings (around 30-35 seedlings) were transferred into plastic containers holding half-strength Hoagland’s nutrient solution (pH 5.8) (Hoagland and Snyder 1933) and grown at 28 ±3°C with 14 h photoperiod. Every 2 days interval Hoagland solution was renewed, seven days old seedlings were treated with 10µM (moderate) and 50µM (high) Cadmium chloride (CdCl_2_) in the presence and absence of 5µM Sodium selenite (Na_2_SeO_3_), used as a source of Selenium (Se). These environmentally related CdCl_2_ concentrations were selected from the previous report (Xie et al. 2015; Barman et al. 2020). Based on our preliminary work 5µM Se was selected as a moderate concentration from three different Se concentrations (2.5µM, 5µM and10µM). 5µM Se was found to be optimally effective to enhance growth and photosynthetic pigment contents (Supplementary Fig. S1). Therefore, the experiment was designed with six treatment sets as follows: (i) **C** (0 Cd+0 Se), (ii)**T_1_**(5μM Se), (iii)**T_2_** (10μM CdCl_2_), (iv)**T_3_**(5μM Se + 10μM CdCl_2_), (v) **T_4_**(50μM CdCl_2_), (vi)**T_5_**(5μM Se + 50μM CdCl_2_). After 14 days seedlings were harvested and used for the different experiments.

### 2.2. Plant growth analysis

After 14 days phenotypic appearance of both rice seedlings were documented. At the same time plant height in terms of root, shoot length and dry biomass were recorded. Plant relative water content (RWC) was measured as described before (Barr and Weatherley 1962). Briefly, fresh weight (FW) of the collected plants were measured and then kept into test tubes containing water for 24h. After 24h, these plants were thoroughly blotted and measured for the turgid weight (TW). Then these plants were kept in hot air oven for 72h. After 72h of oven dry, plants dry weight (DW) was measured. Then the plant RWC was determined by the help of following formula.

Relative Water Content (RWC)% = [(FW)-(DW)/(TW)-(DW)]×100.

### 2.3. Sulfur metabolic enzymes activity

The activity of enzyme ATP-sulfurylase (ATP-S; EC 2.7.7.4) was measured according to (Lappartient and Touraine 1996). 100mg root and shoot samples from both rice cultivars were homogenized in 2 ml of 20 mM Tris -HCl (pH 8.0) buffer consisting 10 mM Na_2_EDTA, 2 mM DTT, and 0.01 g ml^−1^ PVP and were centrifuged at 12,000 rpm for 15 min. Then 0.1 ml supernatant, 7 mM MgCl_2_, 5 mM Na_2_MoO_4_, 2 mM Na_2_ATP, and 0.032 U ml^−1^ of sulfate-free inorganic pyrophosphate were used as assay mixture. Another set of assay mixture was prepared in similar process without having Na_2_MoO_4_. Finally, the absorbance was taken at 340nm by spectrophotometer.

O-acetylserine (thiol)lyase (OAS-TL; EC 4.2.99.8) enzyme was quantified as according to (Heeg et al. 2008). 100mg tissue was grounded 50 mM potassium phosphate buffer (pH 7.5) and centrifuged at 12,000 rpm for 10 min. Then the reaction was initiated by adding 5mM Na_2_S and 12.5mM O-acetyl serine, 1/5 volumes of 7.5% (w/v) TCA, 200 µl acid-ninhydrin reagent and incubated at 100°C for 5 min. Finally, the absorbance was noted at 560 nm.

Cysteine was measured according to (Gaitonde et al. 1967). 250mg root and shoot tissue was homogenized by 5% (w/v) 1ml ice-cold perchloric acid and cold centrifuged at 2800g for 30min. Reaction mixture containing 0.5ml of extract, 1 ml acid ninhydrin reagent and were measured at 580nm.

### 2.4. Non-enzymatic antioxidants activity

Reduced glutathione (GSH), oxidized glutathione (GSSG), and total glutathione contents were determined from treated and control samples of both the cultivars according to Anderson (1985). Briefly, root and shoot tissues were homogenized in 3ml of 5% sulfosalicylic acid and followed by centrifugation at 10,000 rpm (4^0^ C) for 10 min. Total 1ml assay mixture containing 0.5ml crude extract, 0.1M potassium phosphate buffer (pH 7.0), 0.5mM EDTA, and 50µl of 3mM dithio-bis-2 nitrobenzoic acid (DTNB). After 5 min incubation, GSH contents were noted at 412nm. Total glutathione content was determined by adding 100µl of 0.4mM NADPH and 2µl glutathione reductase (GR) to the same reaction and allowed to run after 20 min incubation at dark condition. The amount of GSSG was calculated by the following formula GSSG = (total glutathione) - (reduced GSH content). Total non-protein thiol (NP-SH) and theoretical determination of phytochelatin contents (PCs) were measured by (Del Longo et al. 1993) and (Bhargava et al. 2005) respectively. For NP-SH content 100µl crude extract was added 0.5ml 100mM potassium phosphate buffer (pH 7.0), 0.5mM EDTA, 0.5ml of 1mM DTNB. The reaction mixture was incubated 15min and absorbance was taken at 412nm in UV-Visible spectrophotometer (Shimadzu, Japan). The total amount of NP-SH was determined from the standard calibration of cysteine. Phytochelatin content was measured by the following formula PC = (NPSH)-(GSH).

### 2.5. Study of morpho-anatomical changes by Scanning Electron Microscopy (SEM) imaging

External morphology of root tips and stomata were documented by Scanning Electron Microscopy (SEM). For the preparation of transverse section (T.S), root tips were collected from both the control and treated sets and transverse sections were made by razor blade as mentioned earlier (Guha et al. 2018). Then only the thin sections from all experimental sets were dehydrated in a graded ethanol series (30%, 50%, 70%, and 90%) for 20 min. In case of stomata, leaf samples were cut into small pieces (approximately 0.5cm length). Finally, root T.S, and stomata were photographed by Scanning Electron Microscopy (ZEISS EVO-MA 10; Carl Zeiss Pvt. Ltd. Oberkochen). Simultaneously, Energy Dispersive X-Ray Analysis (EDX) was also performed to determine the nature of deposition found on the root surface.

### 2.6. Detection of *in situ* localization of Cd by Fluorescence microscopy

Approximately, 1cm root tips were stained with Leadmium AM green dye for 1.5h in dark condition followed by thorough washing with 0.85% NaCl solution. Then these samples were visualized by fluorescence microscopy (Olympus, Japan, 60X) with 490/520nm excitation and emission wavelength respectively as mentioned by (Li et al. 2012).

### 2.7. Study of transporters gene expression

RNA was extracted from control and treated seedlings by RNA isolation kit (HiMedia) as mentioned by the manufacturer’s instruction. The purity and the concentration of isolated RNA was quantified by Nano Drop spectrophotometer. After that, iScript cDNA kit was used for first strand cDNA synthesis. Primer sequence for *OsIRT1, OsZIP1, OsMTP8, OsCOPT3, OsSULTR1.1,* and reference gene *Os18SrRNA* were selected by Primer3 using FASTA sequences (the sequence of primers are mentioned in Supplementary Table. S1). Finally, the transporters gene expression was performed by RT-PCR using iTaq (Biorad) and calculated by using the 2^−ΔΔCT^ formula.

### 2.8. Pot Experiment

Seasonal pot experiments were conducted from November to March (Annual temperature range: 25°C-35°C, annual precipitation: ≤ 375 mm, humidity: 60%, sunlight/light photoperiod range: 11h-13h) in Agricultural Experimental Farm (88°26.164′E and 22°22.526′N), University of Calcutta, Baruipur, South 24 Parganas, West Bengal, India. 5 kg of surface (0-20 cm) soil was collected from rice field and added to each pot. The surface sterilization and germination process of rice seeds were done in the same way as mentioned earlier. On the 21^st^ day, 4 seedlings (equal size and length) were transplanted into these pot soil and subjected to Se and Cd treatments. During the Se and Cd treatments, we made aqueous solutions of 10μMCdCl_2_, 50μMCdCl_2_ and 5μMNa_2_SeO_3_ and applied accordingly to the pot soil in such a manner, that at least 2 cm of standing water remained over the soil. The six treatment groups were maintained separately as mentioned earlier like (i) **CP** (0 Cd+0 Se), (ii)**P_1_**(5μM Se), (iii)**P_2_** (10μM CdCl_2_), (iv)**P_3_** (5μM Se + 10μM CdCl_2_), (v) **P_4_**(50μM CdCl_2_), (vi)**P_5_** (5μM Se + 50μM CdCl_2_). Experiments were conducted in completely randomized design (CRD). Then the plants were maintained in poly house with watering at regular intervals. Agronomic parameters including plant height, leaf length, and width were recorded at the vegetative and mature stage.

### 2.9. Soil analysis

Soil was sampled from each pot and pH was measured. Different soil Cd fractions were measured through a selective sequential extraction method by the standard protocol of (Yong et al. 1993). Each fraction was collected and Cd contents were calculated according to (Mohamadiun et al. 2018).

### 2.10. Rice grain, root and shoot element investigation by ICP-AES

After maturity, harvested grain, root, and shoot samples were ground into powder and digested by tri-acid mixture of HNO_3_: HCl: HClO_4_ (4:2:1) with 120°C temperature in a block digestion chamber. After that, samples were kept overnight for cooling and element (such as Fe, Mn, Cu, Zn, S, Cd, Se) content was measured by ICP-AES.

### 2.11. Estimation of rice starch, soluble sugar, protein and thiamine (vitamin B1) content

Starch and soluble sugar content of rice seeds were estimated by the standard protocol as given by (McCready et al. 1950) and (Dubois et al. 1956) respectively. As per the protocol, 10mg dehusked seeds were homogenized in 0.4 ml 80% ethanol and allowed to kept in boiling water bath at 95°C for 10 min. After boiling, pellet was collected by the centrifugation of that sample at 2000 rpm for 20min. The supernatant was kept for soluble sugar analysis. For starch estimation, collected pellet was suspended in 0.65 ml of 50% perchloric acid and kept at 8°C for 24h. After incubation samples were centrifuged at 5000 rpm for 5 min and read at 620 nm. For soluble sugar estimation, 200 μl supernatant was added with 4 ml of anthrone reagent and allowed to incubate in a boiling water bath for 10 min at 100°C. Finally, the absorbance was noted at 620nm. Thiamine and protein content were estimated from post harvested rice seeds from each treatment set as mentioned by (Bhattacharyya and Roy 2018) and (Lowry et al. 1991) respectively.

### 2.12. Estimation of daily intake of Cd, Se and calculation of associated risk factor

Estimated Daily Intake (EDI) of Cd through rice and its associated Cancer Risk (CR) were calculated by the following formula as mentioned before (Naseri et al. 2018) and (Nduka et al. 2019) respectively.

Estimated daily intake (EDI) = (C × Cons)/W Cancer Risk (CR) = EDI × CSF

Where C = Cd content present in rice (mg/kg), Cons = amount of rice consumption by an individual (500g/day), W = average body weight (60 kg for adult). CSF means Cancer Slope Factor = 0.38 for Cd (Nduka et al. 2019). At the same time Se intake rate was also calculated.

### 2.13. Statistical analysis

Statistical analysis was done by using the SPSS software ver.21. Tukey test was performed to determine the significant differences between the treatment and control sets at the level of (p ≤ 0.05).

## 3. Results

### 3.1. Se supplementation improved plant growth

Plant phenotypic appearance and growth were documented in figure 1. This finding showed that Cd stress reduced the growth of both of the rice seedlings as a result root length, shoot length and biomass were reduced. Root inhibition was more pronounced in the T_4_ set of Jamini compared to the rest of the cultivar. But Se supplementation positively regulates the plant growth when combined with Cd. As the growth parameters improved by Se supplementation so the plants relative water content (RWC) was also enhanced in Se exposed sets.

**Fig. 1.**
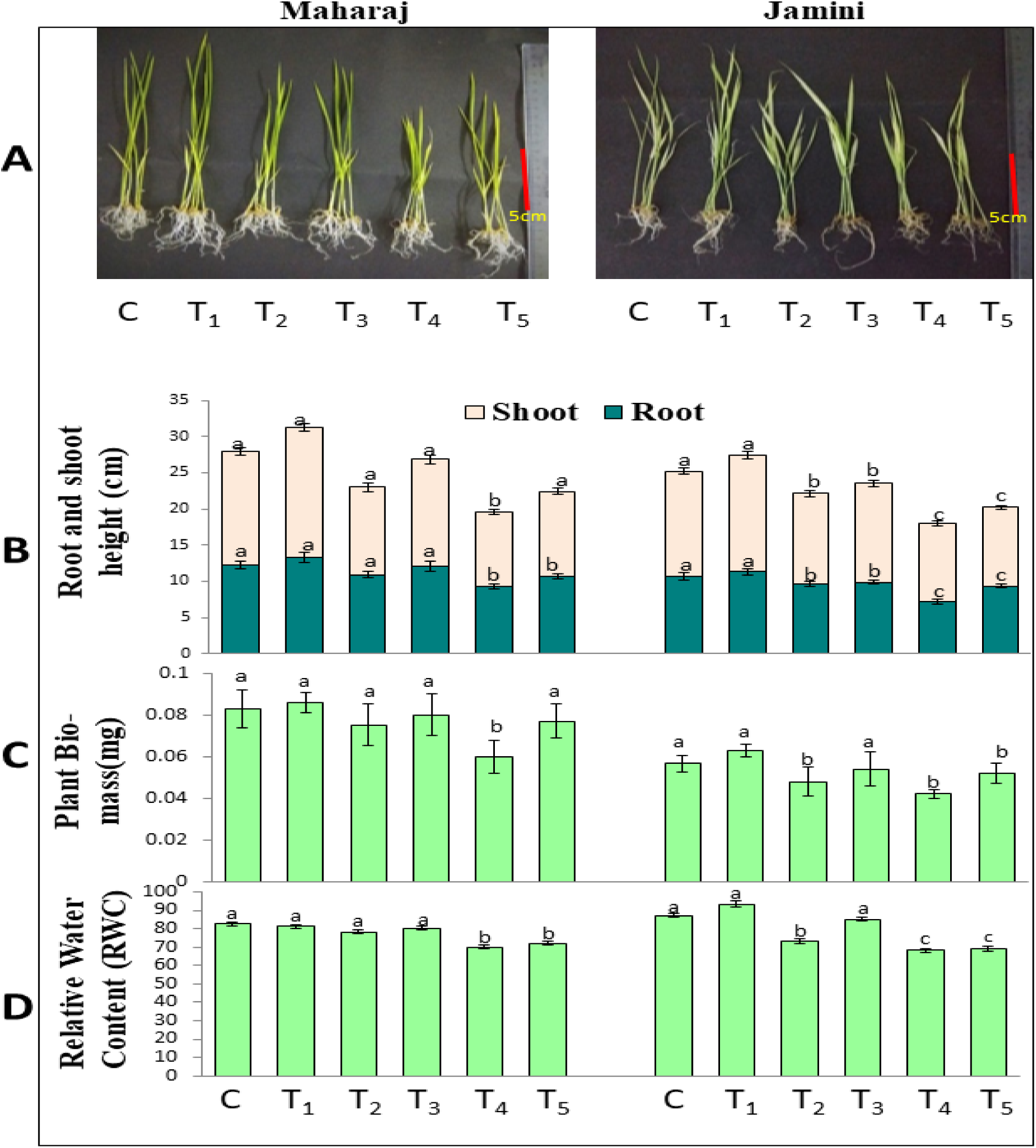
(A) Phenotypic appearance of both cultivars (scale bar =5cm). (B) Plant height (C) Plant biomass and (D) Relative water content (RWC). The presented data were mean value with ± SE of five independent replicates (n =5). The bar graphs marked with same letters indicating these values are not considered as significantly different on the basis of Tukey’s HSD analysis where p ≤ 0.05. [**C** (0 Cd+0 Se), **T_1_**(5μM Se), **T_2_** (10μM CdCl_2_), **T_3_** (5μM Se + 10μM CdCl_2_), **T_4_** (50μM CdCl_2_), **T_5_** (5μM Se + 50μM CdCl_2_)].

**Table. 1.** Estimated daily intake (EDI) of Cd and associated cancer risk. [**P_2_**(10μM CdCl_2_), **P_3_** (5μM Se + 10μM CdCl_2_), **P_4_** (50μM CdCl_2_), **P_5_** (5μM Se + 50μM CdCl_2_)].

| Sample | Treatment Set | Cd content detected in rice (mg/kg) | EDI for 500g rice (µg/kg BW/day) | EDI for 1kg rice (µg/kg BW/day) | Cancer Risk for 500g rice (EDI×0.38) | Cancer Risk for 1kg rice (EDI×0.38) |
| --- | --- | --- | --- | --- | --- | --- |
| **Maharaj** |  |  |  |  |  |  |
|  | **P2** | 0.025 | 0.208a | 0.416a | 0.079a | 0.158a |
|  | **P3** | 0 | 0 | 0 | 0 | 0 |
|  | **P4** | 0.075 | 0.625b | 1.25b | 0.237b | 0.475b |
|  | **P5** | 0.033 | 0.275a | 0.55a | 0.104a | 0.209a |
| **Jamini** |  |  |  |  |  |  |
|  | **P2** | 0 | 0 | 0 | 0 | 0 |
|  | **P3** | 0 | 0 | 0 | 0 | 0 |
|  | **P4** | 0.041 | 0.341a | 0.682a | 0.129a | 0.259a |
|  | **P5** | 0.028 | 0.233b | 0.466b | 0.088b | 0.176b |

### 3.2. Se modulates S metabolism

The activity of ATP-S, OSA-TL and cysteine levels increased in Se treated sets (Fig. 2). Although this increment was not significant compared to control sets. Mostly, the T_4_ sets showed a remarkable decline in S metabolic enzyme’s activity. In T_4_ set of roots, the activity of ATP-S, cysteine, and OSA-TL content were decreased by 20%, 25%, and 17% respectively in Maharaj and 15%, 52%, and 51% respectively in Jamini cultivar (Fig. 2 A-C). But Se induced these enzymes activity when combine with Cd i.e. in T_5_ sets of both cultivars. Where as in shoot tissue ATP-S activity was enhanced by (38 and 40% in Maharaj) and (55 and 36% in Jamini) in T_3_ and T_5_ sets compared to T_2_ and T_4_ sets. Increased cysteine content was observed in root tissue in T_3_ (20 and 22%) and T_5_ (20 and 5%) sets as compared to T_2_ and T_4_ groups of Maharaj and Jamini respectively.

**Fig. 2.**
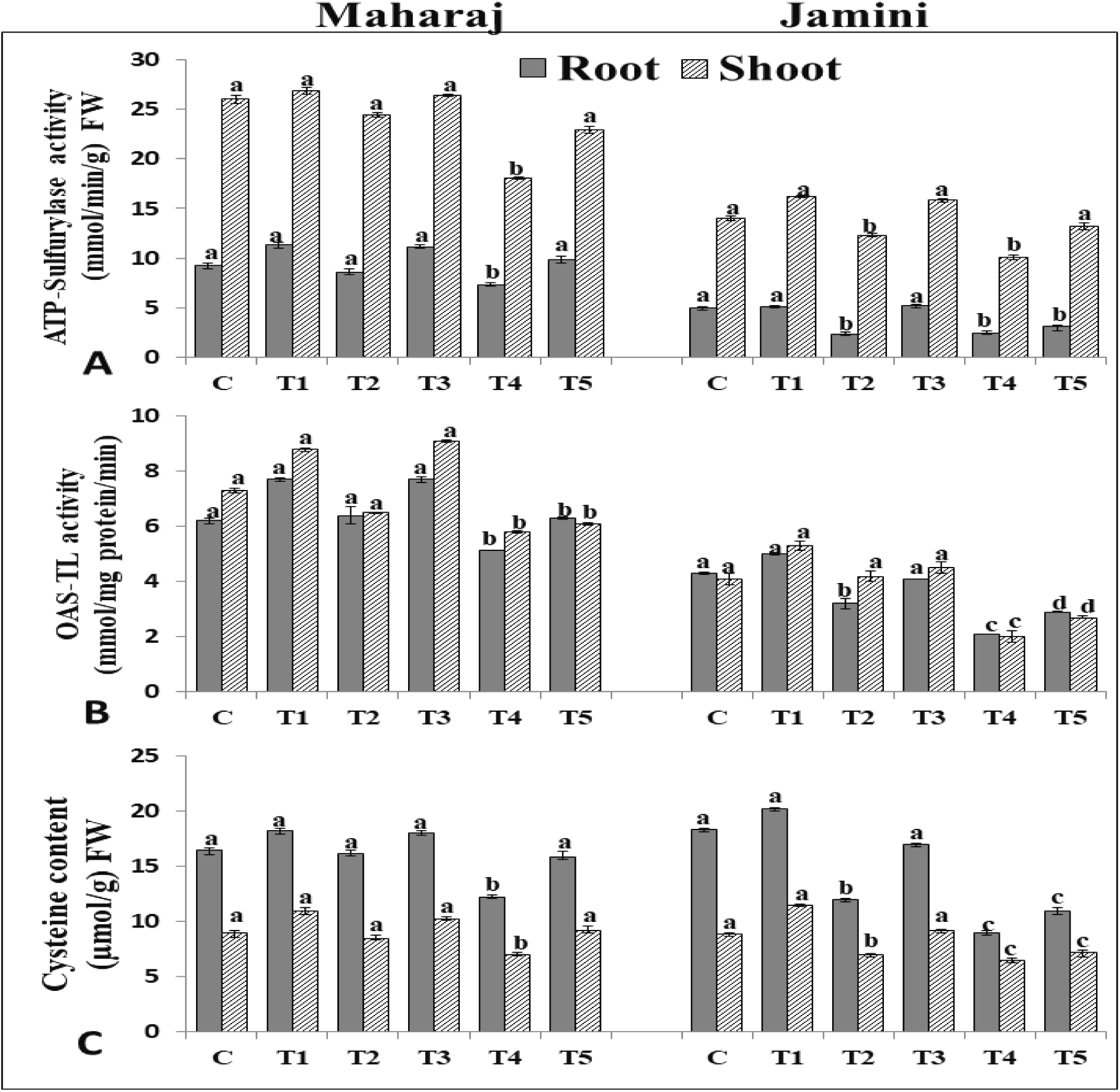
Effect of Se treatment on the activity of Sulfur metabolizing enzymes under Cd stress in 14 days old rice seedlings (A) ATP-sulfurylase activity (B) O-acetyl serine(thiol)lyase (OAS-TL) (C) Cysteine content. The given data are mean of three replicates (n=3) and error bars indicate standard error. The bar graphs containing same letters are not significantly different on the basis of (Tukey’s HSD test at p ≤ 0.05). [**C** (0 Cd+0 Se),**T_1_**(5μM Se),**T_2_**(10μM CdCl_2_), **T_3_** (5μM Se + 10μM CdCl_2_), **T_4_** (50μM CdCl_2_), **T_5_** (5μM Se + 50μM CdCl_2_)].

### 3.3. Se induces sulfur containing thiol compound’s activity

Experimental result indicated that Cd treatment reduced the NP-SH level in the root (16, 41%) and shoot (15, 14%) in 50µM CdCl_2_ treated sets i.e. T_4_ sets in Maharaj and Jamini cultivars respectively (Fig. 3). Exogenous Se treatment slightly enhanced the NP-SH level in Jamini but, Maharaj showed significant increment (136, 112%) and (4, 35%) in T_3_ and T_5_ treated sets compared to only Cd treated groups (T_2_ and T_4_). Increased PC level was found in root tissue accounting to 42 and 63% in T_4_ groups of Maharaj and Jamini respectively. Additionally, Se mediated increase in the level of GSH, GSSG, PCs were recorded in T_3_ and T_5_ groups (Fig. 3B-D).

**Fig. 3.**
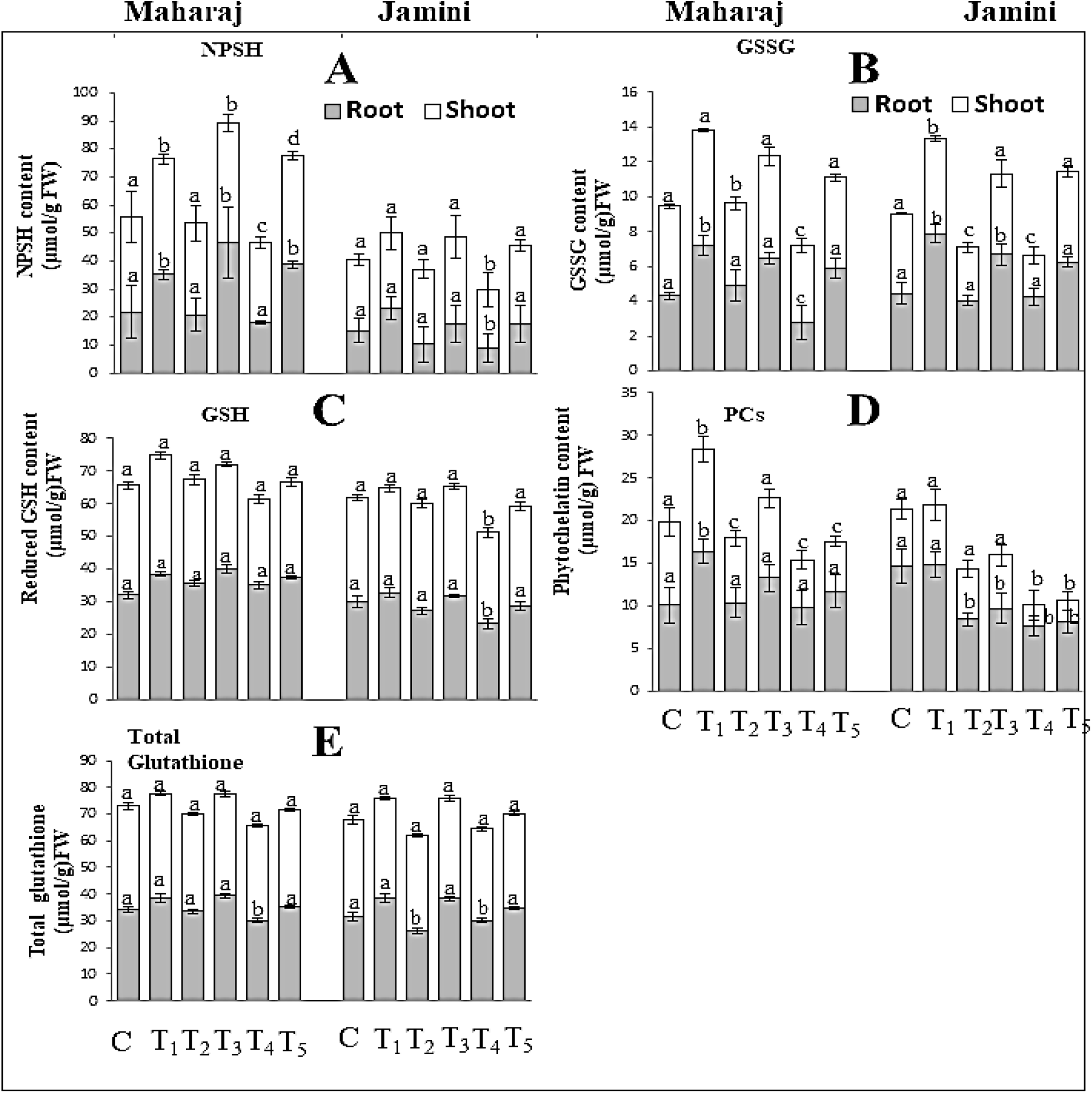
Effect of Se treatment on non-enzymatic antioxidant profiles (A) NPSH content (B) GSSG content (C) Reduced glutathione (GSH) activity (D) PCs content (E) Total glutathione content. The given data are mean of three replicates (n=3) and error bars indicate standard error. The bar graphs containing same letters are not significantly different on the basis of (Tukey’s HSD test at p ≤ 0.05). [**C** (0 Cd+0 Se),**T_1_**(5μM Se),**T_2_**(10μM CdCl_2_), **T_3_** (5μM Se + 10μM CdCl_2_), **T_4_** (50μM CdCl_2_), **T_5_** (5μM Se + 50μM CdCl_2_)].

### 3.4. Se supplementation improved Cd-induced morpho-anatomical changes

SEM images revealed the morphology of root tip, internal structure of root and stomatal opening of 14 d old rice seedlings (Fig. 4). Seedlings subjected to higher Cd concentration i.e. T_4_ showed fully closed stomata in both rice cultivars. T_4_ sets, of both the cultivars showed fully closed stomata. However, partially open stomata were observed in T_2_ set of Jamini. Supplementation with Se promoted stomatal opening in Se+Cd groups (T_3_ and T_5_). Normal cellular structure and morphology were visualized in both control and Se supplemented roots but Cd exposure caused tearing and damages in root tip epidermal cells. In comparison to the Maharaj cultivar, Jamini showed greater tearing of epidermal cells and vascular bundle for T_2_, and T_4_ groups (indicated by the yellow arrow) (Fig. 4k and 4q). EDX analysis of root surface indicated that Se application promotes Cd immobilization and restricts Cd on the root outer surface. As a result, increased Cd concentration was found in Se+Cd i.e. (T_5_) treatment sets compared to only Cd groups (T_4_) in both cultivars (Supplementary Fig. S2).

**Fig. 4.**
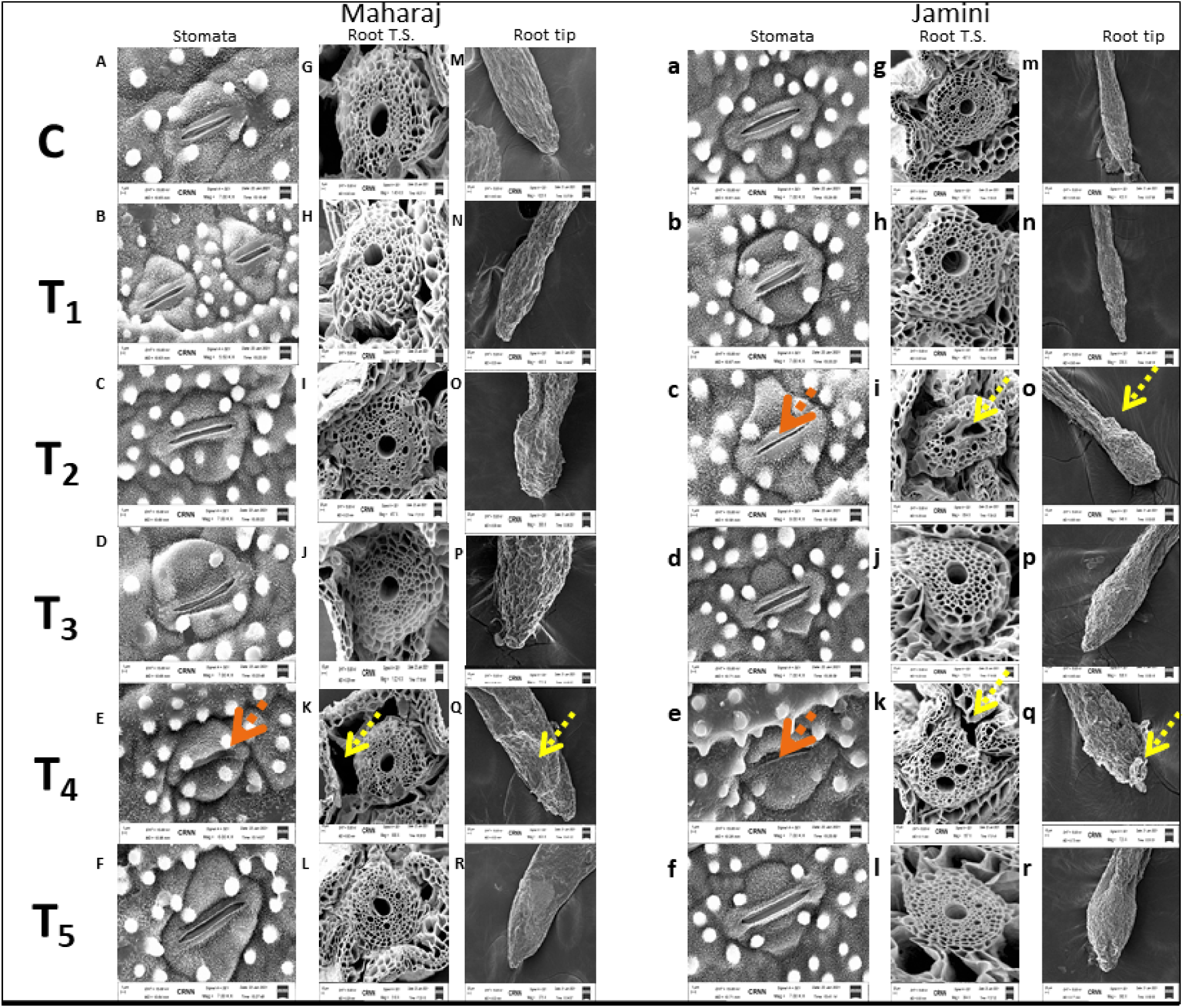
Scanning electron micrograph (SEM) of 14 days old rice seedlings under Se and (Se + Cd) co-treated condition, stomata (A-F) and (a-f), root T.S. (G-L) and (g-l), root tip (M-R) and (m-r) of both Maharaj and Jamini cultivar respectively. Orange arrow indicating fully and partially closed stomata and yellow arrow indicating damaged epidermal cells and root surface. Scale bar =1,5 and 20 µm for stomata, root T.S, and root tip respectively.[**C** (0 Cd+0 Se),**T_1_** (5μM Se),**T_2_**(10μM CdCl_2_), **T_3_** (5μM Se + 10μM CdCl_2_), **T_4_**(50μM CdCl_2_), **T_5_** (5μM Se + 50μM CdCl_2_)].

### 3.5. Se reduced Cd distribution in rice root

The role of Se inhibiting Cd uptake and distribution along rice root was further evaluated by using cadmium specific Leadmium Green AM dye by fluorescence microscopy. Clear and deep green fluorescence throughout the roots (from tips to upper regions) was observed in both the cultivars when roots were treated with only Cd (T_4_ sets). But when seedlings were treated with Se in combination with Cd i.e. (T_5_ sets) the intensity of green fluorescence was much reduced indicating lesser Cd uptake and distribution in the root tissue (Fig. 5).

**Fig. 5.**
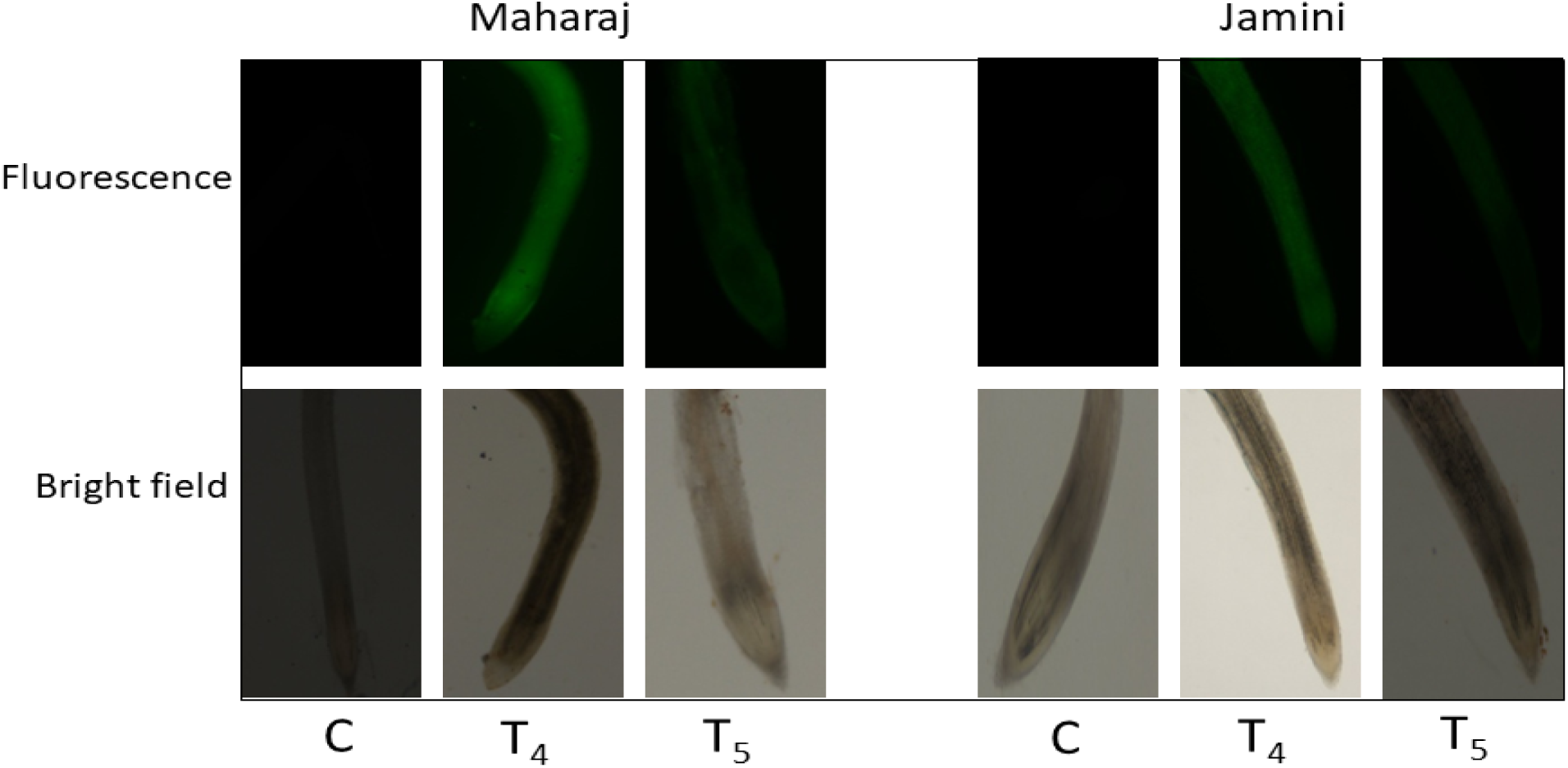
Fluorescence micrographs of rice roots showing *in situ* Cd localization by staining with Leadmium Green AM dye. [C (0 Cd+0 Se), T_4_ (50μM CdCl_2_), T_5_ (5μM Se + 50μM CdCl_2_)].

### 3.6. Expressions of transporter gene under Se and Cd treatments

The transcript level of nutrient transporters was modulated in T_4_ as compared to control. The expression level of *OsIRT1* transporter was up-regulated by 0.72 fold in Maharaj (Fig. 6 A-B). Whereas, T_5_ i.e. Se+Cd co-treated set showed down regulation of *OsIRT1* transporter in both cultivars. A significant up-regulation of *OsMTP8* gene was noticed in shoot tissue of Maharaj (59 fold) and root of Jamini (15 fold) upon Se+Cd treatment compared to control (Fig. 6C-D). Similarly, *OsZIP1*and *OsCOPT3* transporters were also found to be down regulated in Maharaj and Jamini cultivar respectively in T_5_ sets compared to T_4_ set (Fig. 6E-H). Supplementation with Se showed a higher transcript level of the *OsSULTR1.1* gene in both the cultivars of Se and Se+Cd treatment groups when compared to their respective Cd sets (T_4_) (Fig. 6 I-J).

**Fig. 6.**
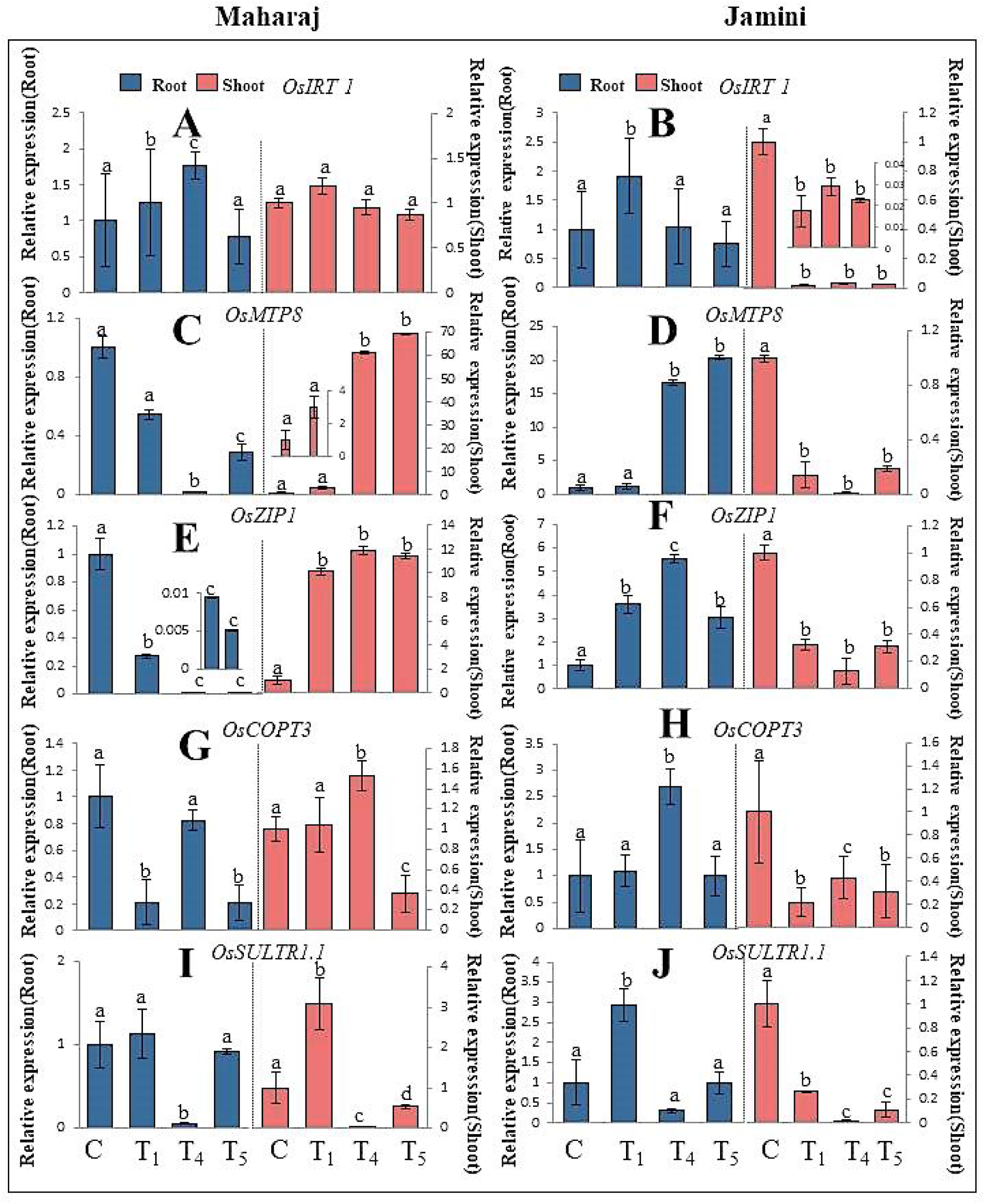
Relative expression analysis of nutrient transporter genes, (A and B) *OsIRT1*, (C and D) *OsMTP8*, (E and F) *OsZIP1*, (G and H) *OsCOPT3*, (I and J) *OsSULTR1.1* transporter in root and shoot of Maharaj and Jamini cultivars by real time PCR (RT-PCR). Data represent the mean of three replicates (n=3) and error bars represent standard error. The bar graphs containing same letters are not significantly different on the basis of (Tukey’s HSD test at p ≤ 0.05). [**C** (0 Cd+0 Se),**T_1_** (5μM Se), **T_4_** (50μM CdCl_2_), **T_5_** (5μM Se + 50μM CdCl_2_)].

### 3.7. Agronomic traits and yield components were enhanced by Se under Cd stress

As shown in (Fig. 7A) retarded growth due to Cd toxicity was observed in both cultivars. The higher concentration of Cd treatment (P_4_) significantly reduced the plant initial height by 18% in Jamini cultivar during the vegetative stage but supplementation with Se rescued this toxic effect in (Se+50µM CdCl_2_) treated set i.e. P_5_, and growth was restored by 12% and showed better phenotypic appearance. During maturation, similar trend was observed in Jamini cultivar whereas better tolerant potential was noticed in Maharaj. Plant biomass was significantly reduced under Cd stress in Jamini by 20 and 35% in P_2_ and P_4_ exposed groups, but no such significant reduction was found in Maharaj. Supplementation with Se enhanced the biomass and different yield attributes including effective tiller, panicle length, yield, grain weight under Cd treatment condition (Fig. 7 C-F and Supplementary Table. S2).

**Fig. 7.**
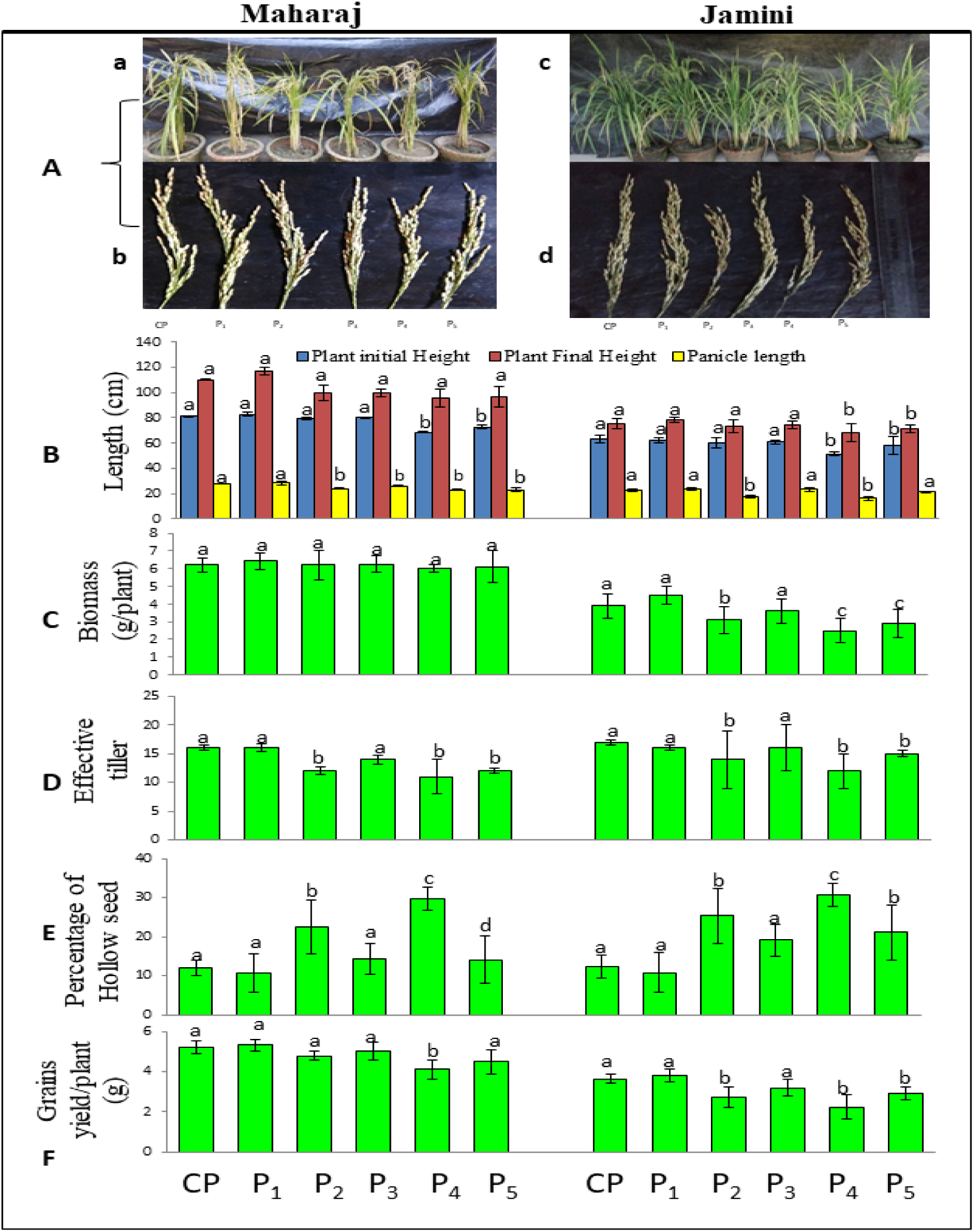
Phenotypic observation, agronomic yield parameters of Se and Se+Cd treated plants. Phenotypic appearance of reproductive stage A (a & c) and panicle A (b & d) of Maharaj and Jamini cultivar respectively. (B) Plant initial, final and panicle length (C) Plant Biomass (D) Effective tiller number (E) Hollow seed% (F) Grain yield/plant. Data represent the mean of ten replicates (n=10) and error bars represent standard error. Means with same letters within each series are not significantly different (Tukey’s HSD multiple comparison at p ≤ 0.05). [**CP** (0 Cd+0 Se), **P_1_** (5μM Se), **P_2_** (10μM CdCl_2_), **P_3_** (5μM Se + 10μM CdCl_2_), **P_4_** (50μM CdCl_2_), **P_5_** (5μM Se + 50μM CdCl_2_)].

### 3.8. Se assisted enhancement of rice grain, root and shoot nutrients, phytochemicals and reduce Cd uptake and cancer risk

Due to Cd toxicity nutrient content was reduced in grains of both cultivars (Fig. 8). Grain Fe content of Maharaj was reduced by 38 and 43% in P_2_ and P_4_ sets and Mn content by 49 and 54%. Whereas Jamini cultivar showed reduced content of Fe (16, 36%) and Mn (32, 44%) in P_2_ and P_4_ sets respectively. Plants treated with only Se enhanced S content by 37 and 30% in Maharaj and Jamini cultivars respectively. Similarly, Se content was also increased in only Se treated sets (Fig. 8F). Cd was detected in P_2,_ P_4_ and P_5_ grains of Maharaj and P_4_ and P_5_ grains of Jamini (Fig. 8G). Supplementation with Se completely reduced grain Cd in P_3_ sets of Maharaj. But when Se was combined with higher concentration of Cd i.e. P_5_ set, it reduced 55 and 40% grain Cd in

**Fig. 8.**
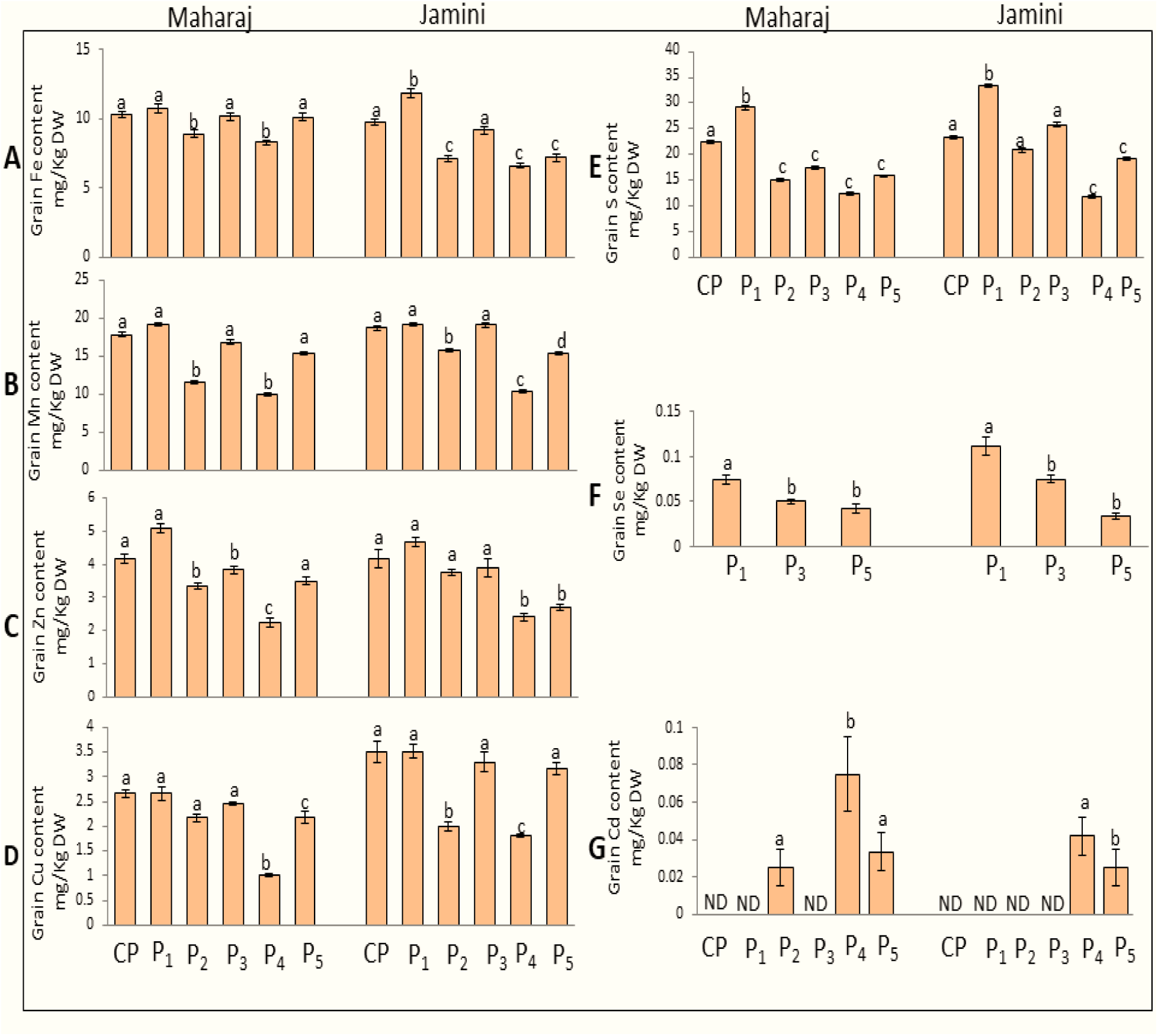
Nutrient and Cd content of harvested rice grains under Cd and (Se+Cd) co-treated sets, (A) Fe content (B) Mn content (C) Zn content (D) Cu content (E) S content (F) Se content (G) Cd content. Data represent the mean of three independent replicates (n=3) and error bars represent standard error. The bar graphs containing same letters are not significantly different on the basis of (Tukey’s HSD test at p ≤ 0.05). (ND means not detected). [**CP=** control plants (0 Cd+0 Se), **P_1_** (5μM Se), **P_2_**(10μM CdCl_2_), **P_3_** (5μM Se + 10μM CdCl_2_), **P_4_** (50μM CdCl_2_), **P_5_** (5μM Se + 50μM CdCl_2_)].

Maharaj and Jamini respectively. Simultaneously, rice root and shoot Fe, Mn, Zn, Cu, S levels were enhanced in (Se+Cd) co-treated sets when compared to only Cd treated sets (Supplementary Figure. S5). Moreover, rice grain’s starch, sugar, thiamine content and protein level were enhanced in Se+Cd treated plants compared to only Cd exposed plants (Supplementary Figure. S4). Importantly, Se supplementation prevents Cd uptake from soil through rice root and shoot tissue. As a result significantly low level of Cd was found in root and shoot sample (i.e. in P_3_ and P_5_ sets) as compared to only Cd exposed sets (P_2_ and P_4_ sets) (Supplementary Table S3).

Bio-concentration factor (BCF) considered as enrichment of toxic metal content in plants, was significantly reduced upon Se treatment in both cultivars. TF i.e. translocation factor (shoot/root Cd content) represents transferability of toxic metal from root to shoot and was also reduced by Se treatment. As Cd was detected in P_2,_ P_4_ and P_5_ grains of Maharaj and in P_4_ and P_5_ grains of Jamini rice variety. So, the estimated daily intake (EDI) of Cd and cancer risk (CR) for both rice consumption was calculated and presented in (Table. 1). On the basis of data (Table. 1) estimated daily intake (EDI) of Cd through Maharaj rice consumption (500g/day) will be 0.208 and 0.625µg/kg BW/day for P_2_ and P_4_ grains. Whereas EDI of Cd for Jamini rice consumption (500g/day) will be 0 and 0.341µg/kg BW/day for P_2_ and P_4_ sets. Interestingly, Se supplementation completely reduced EDI of Cd as well as cancer risk in P_3_ sets of both cultivars. But when Se combined with high concentration of Cd i.e. P_5_ sets then the EDI of Cd will be 0.275 and 0.233µg /kg BW/day in Maharaj and Jamini respectively.

### 3.9. Se showed no health risk due to Se rich rice consumption

Se intake rate through the rice was evaluated and data presented in the Supplementary Table S4. The highest Se intake will be 56µg from 500g rice, if people consume Jamini rice. Whereas low level of Se intake will be 16µg from 500g Jamini rice consumption. Additionally, Se intake rate will vary from (16-56µg) for both of the rice consumption.

### 3.10. Soil Cd bio-availability reduced by Se

Cd and Se+Cd treated soil was analyzed from pots of both the cultivars, and the result is shown in (Supplementary Fig. S3 B-C). Carbonate fraction was significantly reduced by 14 and 18% in P_3_ and P_5_ exposed groups of Maharaj. Whereas slightly higher percentage of carbonate fraction was found in Jamini i.e. 17 and 25% in P_3_ and P_5_ sets. Beside this soil oxide, organic and residual fractions were significantly increased due to Se supplementation.

### 3.11. Soil pH increased by Se

Slightly low soil pH was found in P_4_ treated set compared to the rest of the treatment sets. However, supplementation with Se enhanced the soil pH in P_5_ sets of both the cultivars (Supplementary Fig. S3 A).

## 4. Discussion

Cd contamination is a serious threat to agriculture and human health, so several strategies were advised for remediation, while physico-chemical methods were found to be expensive and associated with secondary pollution, application of manure, lime, biochar, micronutrients, compost, zeolite and bioremediation can be considered as ecofriendly, sustainable and cost-effective way. They regulate the soil physico-chemical properties and reduce Cd bioavailability by immobilizing and subsequent precipitation (Pramanik et al. 2018). Cd can reduce micronutrients in plants, therefore, supplementation by Fe/Mn was found to reduce Cd accumulation in rice by competitive uptake (Sebastian and Prasad 2014). Iron oxide, coated with modified hairs and zeolite, coated with KMnO_4_, can reduce bioavailable Cd content in soil (Ullah et al. 2020). In the present study, the mitigating effect of Se during Cd stress was carried out in detail.

The plausible mechanism of Se-mediated Cd stress amelioration was further evaluated by studying sulfur metabolism. Selenium (Se) is yet to be confirmed as essential nutrient for plants but it is proved that Se has beneficial role in plants (Hasanuzzaman et al. 2020). Although high concentration of Se can be toxic but several studies have shown that low concentration of Se can enhance plant growth, development and yield. At the same time Se can protect plants from different abiotic stress like heavy metal, drought, temperature, salinity etc. (Balal et al. 2016; Pereira et al. 2018). Thus, Se may act as a vital element for plants by altering physiological and biochemical processes (Hasanuzzaman et al. 2020). Since Se (Selenate form) and S are chemically similar (Chauhan et al. 2019). Unlike Selenate, the selenite does not function as chemical analogs of S, and hence they can be considered to be safer in terms of toxicity (Huang et al. 2015). Selenite is taken up by plants via phosphate transporters, aquaporins, and passive diffusion and helps in sulfur assimilation in plants (Sors et al. 2005). We observed that, under Cd stressed condition Se supplementation enhanced key enzymes responsible for S assimilation including ATP-sulfurylase (ATP-S), O-acetyl serine (thiol) lyase (OAS-TL), and cysteine (Cys) content. A similar observation was also mirrored in reports by Khan et al. (2015), where selenite

treatment in Cd stressed plants lead to up-regulation of ATP-S activity. Since a high concentration of Se can help in S assimilation; the enzymes involved in sulfur metabolism were up-regulated (Turakainen 2007). ATP-S is considered as the metabolic entry point to initiate S assimilation that catalyzes the generation of adenosine phosphosulfate (APS) from ATP and sulfate (Phartiyal et al. 2006). The terminal step of S assimilation is catalyzed by OASTL and the formation of Cys from OAS and H_2_S (Liang et al. 2016). Enhanced activities of ATP-S and OASTL also corroborated with increased cellular Cysteine. Cysteine has essential roles in maintaining protein functions and structures since it is the precursor of several sulfur-containing proteins that directly involved in plant defense signaling (Gotor et al. 2015). Cysteine is also responsible for GSH biosynthesis. GSH has many distinct functions including regulation of gene expression related to maintenance of cellular and sub-cellular redox state maintenance and also acts as a reducing cofactor of enzymes responsible for ROS detoxification. In the present study, the glutathione pool was significantly modulated upon co-treatment with Cd and Se. Although in Cd-treated sets the pool of reduced glutathione declined significantly, Se could restore the reduced glutathione levels. The GSH pools are also involved in the synthesis of phytochelatins (Capaldi et al. 2015). Due to enhanced GSH levels in Se+Cd treated sets (T_3_ and T_5_ sets), the level of phytochelatins and NPSH enhanced significantly. Increase in PC and NPSH contents are directly indicating increased chelation of Cd in the vacuole (Rizwan et al. 2016), which helped to scavenge ROS. It is already established that Se promotes sulfur accumulation in plants. But in our study, we have tried to show Se mediated up-regulation of sulfur metabolism in Cd stressed rice plants and improvement of grain micronutrient content. Previously Sun et al. (2021) reported that Se content enhanced through the up-regulation of S metabolism and reduced arsenic content in rice grains. But Se mediated up-regulation of sulfur metabolizing enzyme activity in Cd stressed rice was not reported earlier. Se also helps to restore S metabolism, as observed from the up regulated expression of *OsSULTR1.1* in Se+Cd treated sets, which showed significant down regulation under Cd exposed condition. Another mechanism for remediation of heavy metal stress involves modulation of the expression levels of transporters involved in Cd uptake, translocation from root to shoot, and sequestration in vacuoles. Cd translocation through xylem is a key factor in shoot/grain Cd accumulation, variation in translocation in genotypes confers tolerance to Cd toxicity. CDF/MTP transporters play an important role in this translocation. Since, Cd has no specific transporters, being a divalent cation it can easily enter the root via nutrient transporters and they often compete with micronutrients like Fe, Cu, Zn, Mn etc. which leads to micronutrient deficiency in the plant (Guha et al. 2020). Regulation of these nutrient transporter genes is considered as another mechanism to reduce Cd uptake from the rhizosphere to plants (Shafiq et al. 2019). So, we have studied the expression of the genes, which are mostly responsible for transport of micronutrients (Cu, Fe, Zn and Mn) as well as Cd (when present). Depending on the cultivars, the expression varies and dictates the tolerant/sensitive nature. An earlier study by (Eroglu et al. 2016) revealed that the transporter *OsMTP8* (Metal Tolerance Protein 8), is a tonoplast located protein involved in Mn, Fe and Zn transport and responsible for vacuolar metal sequestration. It was observed in our study that the transcript level of the *OsMTP8* was reduced significantly under Cd exposed condition, resulted reduced nutrient content. Our result is also supported by Ram et al. (2019), who also found low expression of the *OsMTP8* gene under Cd treatment conditions. The expression varied between the two cultivars, in Maharaj the expression was more in shoots, whereas the expression was more in roots of Jamini in Se+Cd co-treatment. This clearly indicated that, along with other nutrients, more Cd may have entered within the plants but, the tolerant cultivar has its own defense mechanism as, Se supplemented with Cd (i.e. T_5_ groups) enhanced vacuolar sequestration and thereby reduced the concentration of cytosolic toxic metal. Whereas, Jamini being the sensitive cultivar, with not so active anti-oxidant defense mechanism, showed low level of expression in shoots/roots compared to Maharaj. Another transporter *OsZIP1*, acting as an efflux transporter, mostly expressed in roots, is mainly responsible for Zn uptake, and also used to transport Cd. Surprisingly we have observed that, in both the cultivars, the expression was downregulated in T_5_ sets, compared to T_4_ sets (showing more Cd and less Zn accumulation in plants). But, in T_5_ sets, the pattern is opposite, indicating that, due to exogenous Se application bioavailability of Cd has been reduced, resulted lesser Cd and more Zn uptake by the plants. At the same time, Se modulates the plant’s anti-oxidative machinery, biochemical and stress response, and also changed the expression patterns of transporters responsible for Cd uptake conferring tolerance to the plants. However, whether these combined effects can prevent the accumulation of Cd in grains need to be evaluated and hence, a pot experiment was conducted. Prolonged Cd treatment can induce plant growth retardation, lesser number of tiller production, and reduction of yield associated parameters like panicle number, hollow seeds, seed number, and grain yield. Since Jamini is a Cd sensitive cultivar, the maximum reduction in yield was observed in the Cd-treated Jamini plants as compared to Maharaj. However, Se supplementation enhanced the plant’s biomass, growth, and effective tillers which were ultimately reflected as yield increments in both Cd sensitive Jamini and Cd tolerant Maharaj.

Upon Se treatment, overall plant metabolism was enhanced in Se and Se+Cd co-treated plants. This enhanced metabolism is generally associated with an increased nutrient uptake. So, the nutrient increment was observed in only Se supplemented plants as well as Se+Cd co-treated plants (i.e. in P_3_ and P_5_ plants). It is also reported by several authors that higher Cd concentration has more negative impact compared to low Cd concentration (Abbas et al. 2018). As a result plants productivity, starch, sugar etc. contents were also reduced at higher Cd concentration. But these parameters were again improved by the Se supplementation in T_5_ sets (Supplementary Fig. S4). In that way Se enhanced nutrient uptake by plants (i.e. in Se+Cd plants) which were minimized in Cd-treated plants. From the pot experiment, it was found that Se has the potential to immobilize the phytoavailable Cd in soil. Liu et al. (2020) also reported that exogenous application of Se reduced phytoavailability of Cd, which lowers the Cd content in rice grains. The solubility and bio-availability of Cd maintain the decreasing sequence i.e. soluble>exchangeable >carbonate-bound>oxide-bound >organic bound >residual (Li et al. 2009). In this study, exchangeable and carbonate fractions of Cd were reduced due to Se application. These two fractions are generally considered as bio-available/phytoavailable Cd because of their high mobility (Dong et al. 2019), and Cd levels in oxide, organic and residual fractions were increased which rendered the Cd unavailable to plants. Thus, Se can reduce phytoavailable Cd because sodium selenite (Na_2_SeO_3_) and related compounds (like selenide, elemental Se, organic Se) might thermodynamically react with Cd and form Se-Cd complex which is less mobile and relatively non-bioavailable (Ren et al. 2020). Detection of *in situ* Cd localization in roots using Leadmium dye, showed green fluorescence throughout the whole roots, both in Maharaj and Jamini in Cd treated roots, but, Se+Cd treated sets showed light green fluorescence only in the vascular regions of the roots, clearly indicating lesser Cd uptake through the root tips. This can be due to sequestration of Cd as Se-Cd compounds on the root surface, as revealed by the EDX analysis. Root surface analysis through EDX also validated the presence of excess Cd, which may have been caused due to immobilization of Cd on the surface. Se application also reduced Cd level in root and shoot tissue of both the rice cultivars. Because Se may combine with Cd and form less mobile (Se-Cd) complex in soil. As a result Cd uptake rate by root and the translocation from root to shoot was observed during (Se+Cd i.e. P_3_ and P_5_ sets). Among different soil parameters like soil temperature, redox potential, soil pH, and soil nutrient contents, soil pH is the most significant factor influencing Cd bio-availability in plants (Roberts 2014). Our result also showed that Se enhanced the soil pH which is also correlated with decreasing Cd bio-availability in Se+Cd treated plants. Enhancement of soil pH and decrease in Cd bio-availability in plants were previously reported in different soil amendments like Se (Huang et al. 2018), Si (Cai et al. 2020), and nZVI (Guha et al. 2020).

Lastly, the effects of Se on grain Cd levels were evaluated in Maharaj and Jamini cultivars. Se completely prevented the accumulation of Cd in P_3_ treated groups of both cultivars. The supplementation of Se reduced Cd translocation as a result higher amount of nutrient elements were detected in root and shoot tissue in Se+Cd exposed plants. Because it was previously, reported by (Yang et al. 2016) and (Chang et al. 2020) that Cd can share common transporters with essential nutrient elements. In our findings as Se prevent Cd translocation so nutrient element levels were also enhanced in root, shoot and grain. Here we also found that 5µM Se was optimum concentration for plant growth compared to 10µM Se because 10µM Se showed plant growth retardation in terms of low level of chlorophyll content (Supplementary Fig. S1). Higher concentration of Se was found to induce phytotoxicity as evident from membrane degradation, chlorosis, and senescence as it intercedes in plant metabolism, leading to reduced plant growth and grain yield (Akbulut and Çakir 2010; Reis et al. 2020). At the same time, high Se also disrupts molybdenum uptake by plants, interrupting molybdopterin biosynthesis, which is required for nitrate reductase activity and nitrogen assimilation from nitrate (Schiavon et al. 2017). So in many ways, higher concentration of Se is detrimental than beneficial for the plant growth. From literature study we also found that 5μM Se enhanced plant growth (Das et al. 2022). Considering our own data and data from earlier workers, we have decided to use 5µM Se for our work throughout, and all the experimental data was found to be corroborating previous findings. We have also measured the growth response under (Se+Cd) exposed condition and results indicated that 5µM Se enhanced growth and ameliorated stress compared to T_4_ set. Moreover, Se treatment enhances the food security for both of the rice. Because as per FAO/WHO (2011) provisional tolerable monthly intake (PTMI) of Cd is 25 μg/kg BW/month. Thus, Se supplementation can ensure safe consumption of Maharaj and Jamini rice grains when they are grown under low and high Cd concentrations (10 and 50µM). Previously Hartwig (2013) reported that Cd intake leads to carcinogenic effects in human body. As our result showed that Se treatment reduce dietary Cd intake. So, it will be assumed that Se supplemented rice reduces the chances of cancer risk.

Furthermore, Se intake results showed that Se consumption ranges from (16-56) µg/d. The recommended dietary amount is 55 µg/d per person, 14 years or above, with a tolerable upper limit of 400 μg/d (Shenkin 2009). Thus we can ensure that the application of selected Se dose during rice cultivation will not be a threat for human health.

Finally, our results proved that Se can play a pivotal role in restoring Cd stress tolerance in Cd sensitive rice cultivar Jamini which was reflected in the enhanced yield potential of Se+Cd treated seedlings. On the other hand, Maharaj was already reported to be Cd tolerant but a Cd accumulator cultivar. Upon Se supplementation, the stress tolerance levels of Maharaj were increased further and most importantly produced Cd free rice (during low concentration of Cd treated condition). So, the cultivation of Maharaj and Jamini rice genotype in moderate levels of Cd contaminated field will have no health risk, if supplemented externally with selenium.

## 5. Conclusion

Cereal crops like rice, wheat, maize are cultivated all over the world. In present time irrigated lands are contaminated by toxic metals like Cadmium (Cd), Lead (Pb), Chromium (Cr), Arsenic (As), which leads to bioaccumulation of heavy metals into the food chain and poses a human health risk. Due to adverse effect of Cd it is needed to cultivate safe and healthy crops for the growing population. This study indicated that Se alleviates Cd induced toxicity in rice through the anti-oxidative properties, up-regulation of sulfur metabolic enzymes, and regulation of Cd transporter gene expressions. Se also regulates the nutrient transporter genes that help nutrient enrichment in grains of both rice cultivars. Further Se also enhance rice yield under Cd stress condition of both rice cultivars and enhanced Se content in rice that is advantageous for human health. Moreover, rice grain micronutrient contents were also increased, thus improving the nutritional value of rice. So, the current study concluded that Cd stress causes depletion of growth, productivity and micronutrient can be restored through the supplementation of Se. Thus Se supplementation may help to develop a new strategy to produce heavy metal resistant and bio-fortified crop, for sustainable rice cultivation on Cd contaminated sites.

## Acknowledgment

Authors would like to acknowledge CAS, Dept. of Botany, University of Calcutta, DST-FIST for infrastructural and instrumentation facilities. Authors also acknowledge I.I.T Bombay for ICP-AES analysis. FB acknowledges University Grants Commission (UGC), Govt. of India for his fellowship. TG acknowledges Council of Scientific and Industrial Research (CSIR), Govt. of India for her fellowship. Financial assistance from Department of Biotechnology, West Bengal, (Memo No-76(Sanc)BT/P/Budget/RD-14/2017) is also gratefully acknowledged.

## Ethical approval

This article does not contain any studies with human participants or animals performed by any of the authors.

## Availability of data and materials

The authors confirm that the data supporting the findings of this study are available within the article.

## Author contribution

Rita Kundu and Falguni Barman—Conceptualization; Falguni Barman and Titir Guha—Data curation; Falguni Barman —Formal analysis; Rita Kundu — Funding acquisition; Rita Kundu and Falguni Barman —Investigation; Falguni Barman —Methodology; Rita Kundu —Project administration; Rita Kundu —Resources; Falguni Barman —Software; Rita Kundu — Supervision; Rita Kundu and Falguni Barman —Validation; Rita Kundu and Falguni Barman — Visualization; Falguni Barman and Titir Guha — Roles/Writing – original draft; Rita Kundu, Falguni Barman and Titir Guha, Writing – review & editing. All authors read and approved the final manuscript.

