## Supplementary data for "Exogenous selenium supplements reduce cadmium accumulation and restores micronutrient content in rice grains"

**Supplementary Figure**

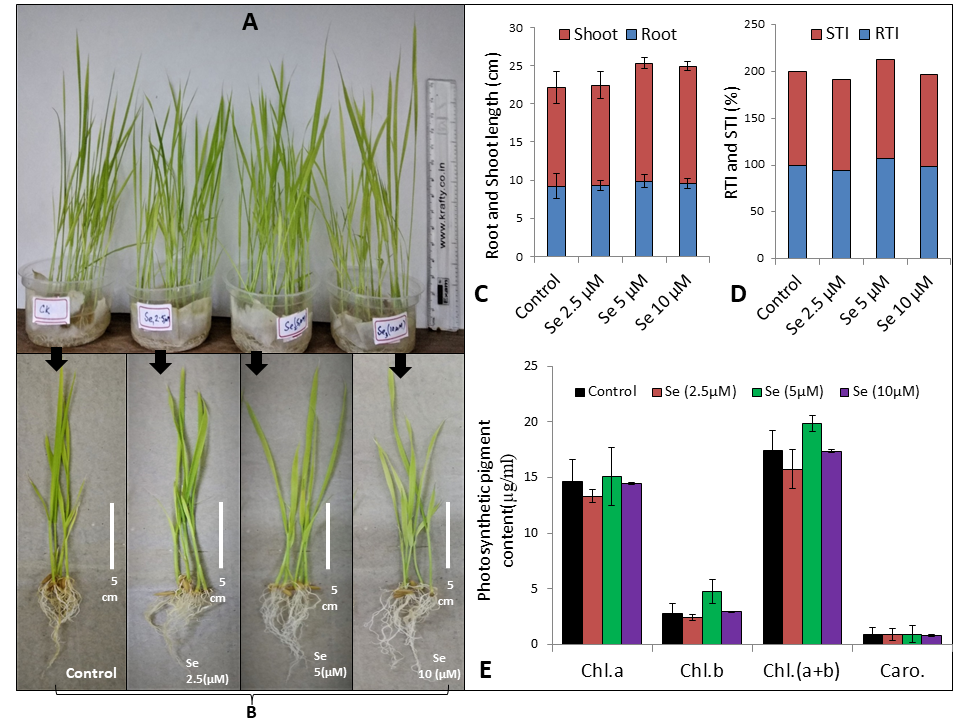

**Supplementary Fig. S1.** (A-B) Phenotypic appearance of 14 days old rice seedlings (Maharaj) under different concentration of Selenium treatments, results obtained after 14 days showed 5µM Se treated seedlings performed better growth. (C) Root and shoot length of different Se treated plants, (D) Root and shoot tolerance index (i.e. RTI and STI) (E) Photosynthetic pigment content.

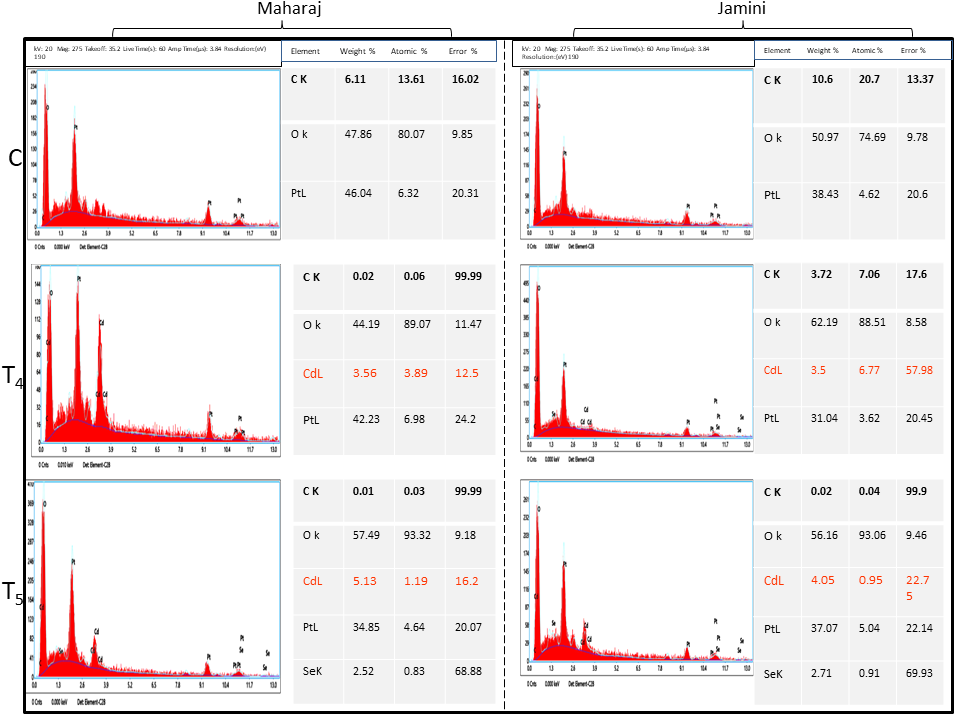

**Supplementary Fig. S2.** EDX analysis of root tip surface showing presence of more Cd in Se-Cd co- treated sets, indicating formation of Se-Cd complex (indicated by red in color) and therefore preventing the Cd accumulation by root. **C** (0 Cd+0 Se), **T_4_** (50μM CdCl_2_), **T_5_** (5μM Se + 50μM CdCl_2_)].

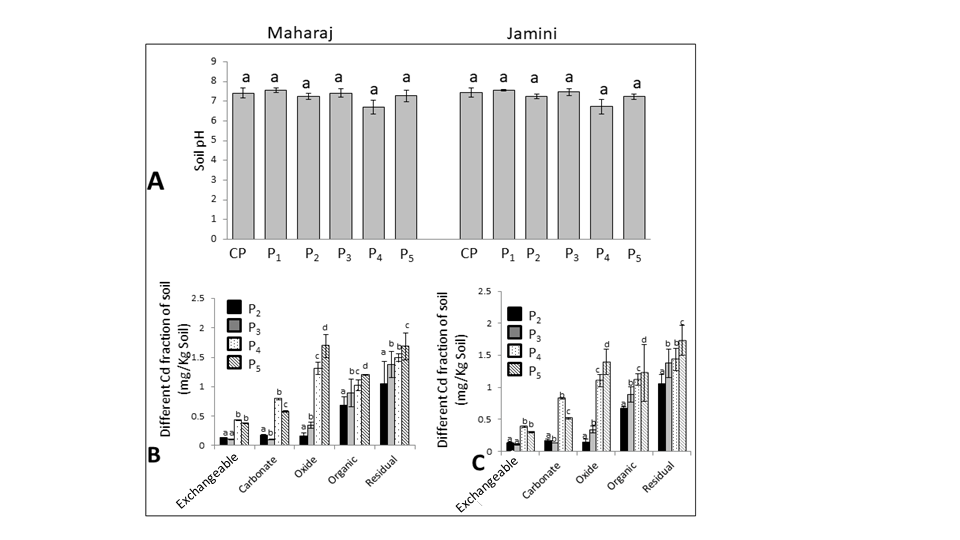

**Supplementary Fig. S3.** (A) Soil pH after Cd and Se treatments of both cultivars and (B-C) represents the effect of Se on Cd levels of soil different Cd fractions after Cd and Se+Cd treatment of Maharaj and Jamini cultivars respectively. Data represent the mean of three replicates (n=3) and error bars represent standard error. Means with same letters within each series are not significantly different (Tukey's HSD multiple comparison at p ≤ 0.05). [**P_2_** (10μM CdCl_2_), **P_3_** (5μM Se + 10μM CdCl_2_), **P_4_** (50μM CdCl_2_), **P_5_** (5μM Se + 50μM CdCl_2_)].

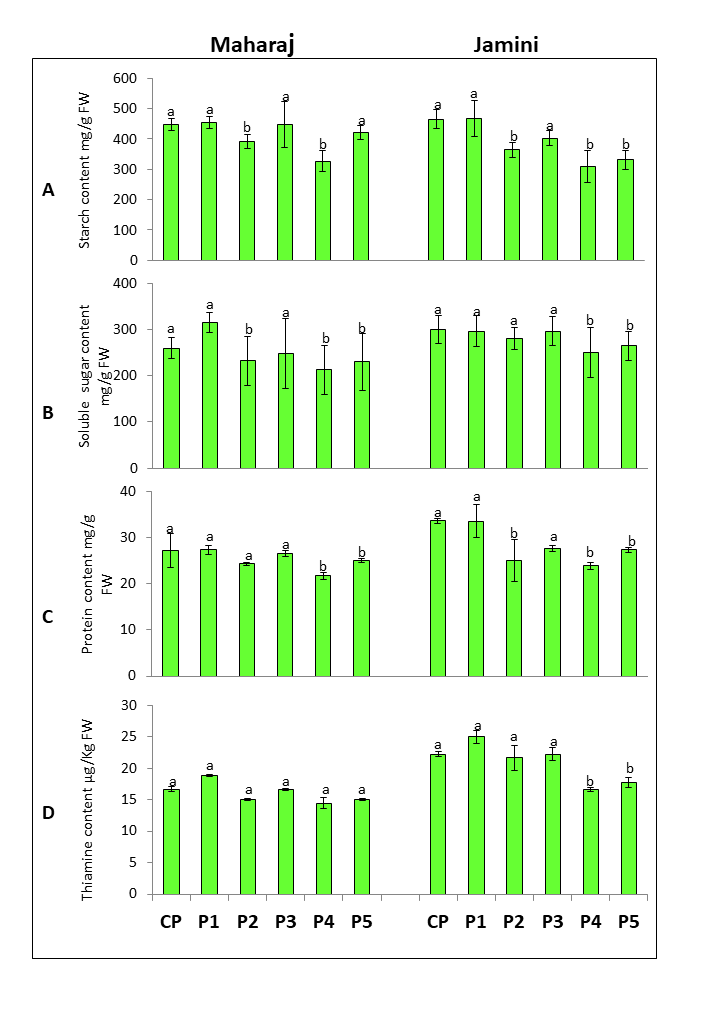

**Supplementary Fig. S4.** Phytochemical analysis of rice grain showing (A). Starch (B). Soluble sugar content (C). Protein content (D). Thiamine content. Data are means ± SD of three independent replica (n=3). [**CP** (0 Cd+0 Se), **P_1_** (5μM Se), **P_2_** (10μM CdCl_2_), **P_3_** (5μM Se + 10μM CdCl_2_), **P_4_** (50μM CdCl_2_), **P_5_** (5μM Se + 50μM CdCl_2_)].

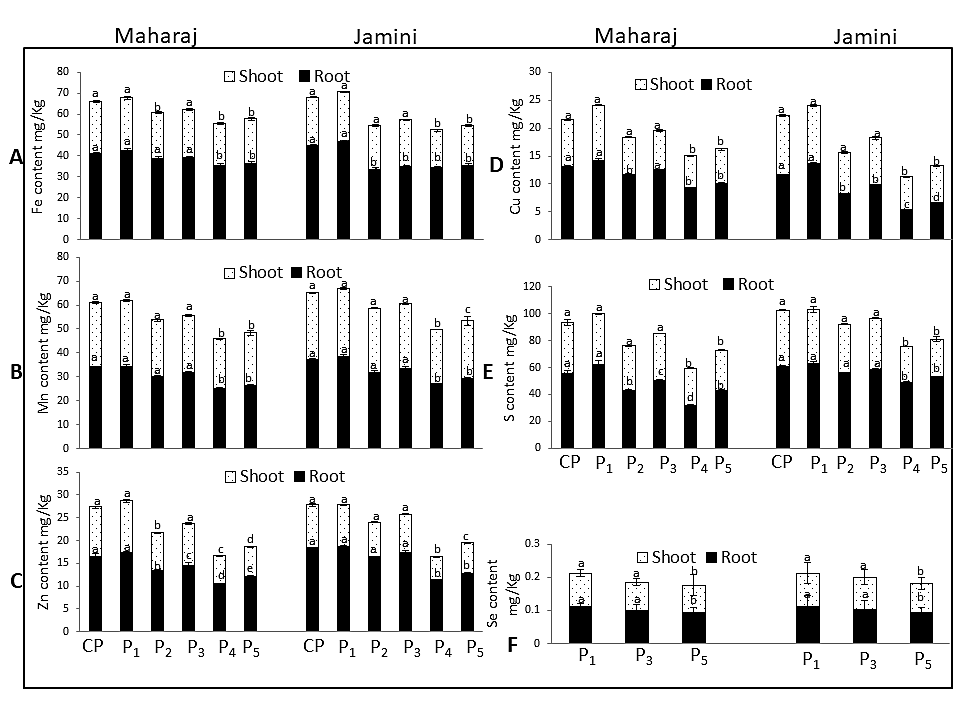

**Supplementary Fig. S5.** Different element content of root and shoot. Data are means ± SD of three independent replica (n=3). [**CP** (0 Cd+0 Se), **P_1_** (5μM Se), **P_2_** (10μM CdCl_2_), **P_3_** (5μM Se + 10μM CdCl_2_), **P_4_** (50μM CdCl_2_), **P_5_** (5μM Se + 50μM CdCl_2_)].

**Supplementary Table**

**Supplementary Table. S1.** Primer sequence for RT-PCR analysis

| Gene | Primer sequences |
| --- | --- |
| *18s* | FW:AACGGCTACCACATCCAAG  RV:CCTCCAATGGATCCTCGTTA |
| *osIRT1* | FW:CGTCTTCTTCTTCTCCACCACGAC  RV:GCAGCTGATGATCGAGTCTGACC |
| *osZIP1* | FW: GACTCCACCTCCACCTTCAA  RW: TGCTCACTGACCTGGTTGTC |
| *osMTP8* | FW: TTACCGGGCTTCTTGATTTG  RW: CATGAACCCGCTGGTAAACT |
| *COPT3* | FW:CGTGCTCCTCGAGTTCCTC  RW:GCGAGCTCTCCCTTCTTGTA |
| *osSULTR1.1* | FW: GTACAAGGACCAGCCGATGT  RW: ATGTCCTGGGGTATGCAGAG |

**Supplementary Table. S2.** Agronomic parameters, values are mean ± SD of ten independent replicates (n=10). [**CP** (0 Cd+0 Se), **P_1_** (5μM Se), **P_2_** (10μM CdCl_2_), **P_3_** (5μM Se + 10μM CdCl_2_), **P_4_** (50μM CdCl_2_), **P_5_** (5μM Se + 50μM CdCl_2_)].

| **Sample** | **Treatment** | **Panicle**  **Weight**  **(g)** | **Panicle**  **/pot** | **100**  **grain**  **Weight (g)** | **Maturation Time ( Days)** |
| --- | --- | --- | --- | --- | --- |
| **Maharaj** | CP | 7.16±0.38a | 17±0.02a | 2.2±0.01a | 135 |
|  | P_1_ | 7.24±0.08a | 22±0.02b | 2.4±0.09a | 134 |
|  | P_2_ | 6.91±0.17a | 16±0.05a | 2±0.04a | 131 |
|  | P_3_ | 7.1±0.11a | 18±0.01a | 2±0.03a | 132 |
|  | P_4_ | 6.05±0.20b | 14±0.21c | 1.7±0.04b | 129 |
|  | P_5_ | 6.91±0.03b | 16±0.08a | 2±0.2b | 131 |
| **Jamini** | CP | 5.21±0.91a | 14±0.05 | 1.9±0.8a | 109 |
|  | P_1_ | 5.52±0.41a | 17±0.09 | 1.9±0.08a | 109 |
|  | P_2_ | 3.91±0.17b | 11±0.11 | 1.6±0.05a | 102 |
|  | P_3_ | 4.02±0.14c | 12±0.33 | 1.8±0.04a | 106 |
|  | P_4_ | 3.25±0.24d | 8±0.08 | 0.8±0.01a | 100 |
|  | P_5_ | 3.58±0.09e | 10±0.02 | 1.2±0.11a | 105 |

**Supplementary Table. S3.** Total Cd content in root and shoot of both rice cultivars. Data represent the mean of three independent replicates (n = 3). Means with same letters are not significantly different (Tukey's HSD multiple comparison at p ≤ 0.05). [**P_2_** (10μM CdCl_2_), **P_3_** (5μM Se + 10μM CdCl_2_), **P_4_** (50μM CdCl_2_), **P_5_** (5μM Se + 50μM CdCl_2_)].

| **Rice Cultivar** | **Treatment**  **Sets** | **Cd content** | |
| --- | --- | --- | --- |
|  |  | **Root (mg/Kg)** | **Shoot (mg/Kg)** |
| Maharaj | P_2_ | 0.86±0.04a | 0.73±0.01a |
|  | P_3_ | 0.19±0.02b | 0.12±0.016b |
|  | P_4_ | 1.43±0.02c | 1.23±0.07c |
|  | P_5_ | 1.19±0.03c | 0.98±0.08c |
| Jamini | P_2_ | 0.75±0.04a | 0.60±0.02a |
|  | P_3_ | 0.16±0.01b | 0.13±0.01b |
|  | P_4_ | 1.39±0.09c | 1.3±0.01c |
|  | P_5_ | 1.06±0.03d | 1.01±0.01d |

**Supplementary Table. S4.** Se intake through dietary rice consumption.

| Treatment sets | Se content in grain | | Dietary intake | |
| --- | --- | --- | --- | --- |
|  | Maharaj (mg/kg) | Jamini (mg/kg) | Maharaj (mg/500gm rice consumed) | Jamini  (mg/500gm rice consumed) |
| P1 | 0.075 | 0.112 | 0.037 | 0.056 |
| P3 | 0.05 | 0.075 | 0.025 | 0.037 |
| P5 | 0.04 | 0.033 | 0.02 | 0.016 |
